# Translation and Transforming Activity of a Circular RNA from Human Papillomavirus

**DOI:** 10.1101/600056

**Authors:** Jiawei Zhao, Eunice E. Lee, Jiwoong Kim, Bahir Chamseddin, Rong Yang, Yang Xie, Xiaowei Zhan, Richard C. Wang

**Author notes:** These authors contributed equally (Jiawei Zhao, Eunice E. Lee, Jiwoong Kim). Corresponding author: Richard C. Wang, MD PhD.

## Abstract

Bioinformatics and in vitro studies have revealed that single-stranded circular RNAs (circRNAs), generated through ‘backsplicing,’ occur more extensively than initially appreciated. While the functions of most circRNAs are unknown, binding of microRNAs (miRNA), regulation of splicing and transcription, and translation into proteins have all been demonstrated for specific circRNAs. Virally-derived circRNAs have recently been described in gamma-herpesviruses. Here, we report that oncogenic human papillomaviruses (HPV) generate circRNAs, including ones which encompass the entire coding region of the E7 oncogene (circE7). HPV16 circE7 can be detected by both inverse RT-PCR and Northern blots of HPV16-transformed cell lines. CircE7 is N^6^-methyladenosine (m^6^A) modified, preferentially localized to the cytoplasm, and can be translated to produce E7 oncoprotein. Specific disruption of circE7 in CaSki cervical carcinoma cells decreased E7 protein levels, inhibited cell proliferation, and inhibited the ability of the cells to form colonies in soft agar. Analysis of TCGA RNA-seq data demonstrates that HPV-positive cancers have abundant circE7 RNAs. These results provide evidence that virally-derived, protein-encoding circular RNAs have biologically important functions with relevance to the transforming properties of HPV.

## Introduction

Human papillomaviruses (HPVs) are small, double-stranded DNA viruses that infect stratified epithelia. While the majority of HPV infections are asymptomatic or cause benign warty growths, a subset of ‘high-risk’ HPVs can promote the development of cervical, oropharyngeal, anal, vulvovaginal, and penile cancers. The recognition of the critical role of HPV infection in the pathogenesis of ∼5% of human cancers has spurred the development of vaccines that have the potential to decrease the burden HPV-driven malignancies^1^. Despite this progress, our understanding of how high-risk HPVs progress from latent infections to incurable cancers remains incomplete.

One strategy employed by HPV to regulate its life cycle is through alternative splicing of its relatively small number of transcripts^2^. In particular, extensive splicing of the early region of high-risk HPV appears to be critical for its tumorigenic properties. The E6 and E7 oncoproteins are transcribed on a bicistronic mRNA, and most E7 oncoprotein translation occurs from a truncated E6 transcript (E6*I) through a mechanism involving translation reinitiation^3, 4^. However, mutations abolishing the 5’ splice donor in the E6 intron do not completely abolish E7 oncoprotein expression suggesting that alternative, uncharacterized transcripts might also contribute to E7 translation^4, 5^. In addition to splicing, many viruses employ non-coding RNAs to promote fitness. Non-coding RNAs, like adenovirus VA and Epstein-Barr Virus (EBV) EBER, prevent the activation of innate immune responses^6^. Similarly, miRNA, like those encoded by EBV and the SV40 polyomavirus, have also been shown to limit activation of the host immune response through targeting of both host and viral targets^7, 8^.

The occurrence of covalently closed single-stranded RNAs (circRNAs) was initially thought to be limited to viroids and rare splicing events from uncommon loci^9^. However, with the recent realization that circRNAs are abundant, interest in this class of RNA molecules has increased. The best characterized circRNAs function as microRNA sponges. For example, circHIPK3, a circRNA derived from the second exon of *HIPK3*, binds multiple miRNAs, including miR-124, to promote the growth of cancer cells in vitro^10^. However, recent studies have suggested additional roles for circRNAs, including the ability of some circRNAs to encode for proteins. Because the translation of circRNAs appears to be markedly lower than that of 5’, 7-methylguanosine capped and polyadenylated transcripts, the biological relevance of circRNA-driven protein production remains unclear. Recent studies have also revealed that EBV and Kaposi Sarcoma Virus (KSHV) generate a diverse menagerie of circRNA^11, 12^. The functions of these viral circRNAs remain uncertain.

We report the discovery of circRNAs from high-risk HPV. Characterization of the abundant HPV16 circE7 revealed that it can be translated through cap-independent mechanisms. HPV-derived circE7 is abundant in cervical and head and neck cancers in The Cancer Genome Atlas (TCGA), and HPV16 circE7 is essential for the transformed growth of CaSki cervical carcinoma cells.

## Results

To screen for the presence of circRNA in HPV, we developed a pipeline to detect and visualize backsplice junctions from viral genomes (vircircRNA). To ensure accurate identification of all backsplices, circular viral genomes were concatenated and then utilized as the reference genome for the pipeline (Fig. S1a-b). We selected ten HPV subtypes (Fig. S1c) as reference genomes to screen against publicly deposited RNA-seq datasets (Fig. S1d). We identified 12 projects with RNA-seq data from HPV-infected tissues. Despite the fact that most samples were not optimized for circular RNA sequencing through RNase R treatment or ribosomal RNA depletion, we identified 27 samples with multiple reads mapping to putative backsplice junctions (Fig. S1d). In our initial screen, backsplice reads were identified from HPV16 and HPV35 (Fig. 1a, S1e). Specific HPV16 circRNAs were notable because their abundance was comparable to spliced linear mRNAs (Fig. 1a-b). The majority of backsplice reads (>93% of total HPV reads, 72% of HPV16- or HPV35-positive samples) were generated from the head to tail joining of established linear splice sites downstream in E1 (nt 16) to one upstream in E6 (nt 7451) (Fig. 1a-c, S1e). This backsplicing is predicted to form a 472nt circular RNA, which contains the entire open reading frame of E7 (Fig. 1c). Due to the established importance of the E7 oncogene and the abundance of HPV16 circE7, we focused on this putative circRNA for additional analyses. We tested for the presence of backsplicing by inverse PCR using three cancer cell lines in which HPV16 has been shown to be stably integrated (CaSki and SiHa cervical cancer, UPCI-SCC154 tongue squamous cell cancer). RNase R is an exoribonuclease that specifically degrades linear, but not circular or lariat, RNAs. While a linear region of HPV16 E6/E7 was markedly decreased in abundance after RNase R treatment, the circE7 junction detected in all 3 cell lines before RNase R was enriched after treatment (Fig. 1d). Sanger sequencing of the amplified inverse PCR product from all three cell lines confirmed that the circE7 backsplice represented a true splice site rather than an intron lariat (Fig. 1e), which frequently contains untemplated nucleotides across the branch point^13^. HPV16 circE7 could be identified in the cancer cell lines by Northern blot as an RNase R resistant band that migrated more slowly than its predicted size due to its circular structure^14^ (Fig. 1f). Given the prevalence of HPV18 in HPV-induced cancers, we also tested 3 HPV18+ cell lines for the presence of a similar circular RNA (HPV18 circE7) (Fig. S2a). However, an analogous HPV18 circE7 could not be detected by either RT-PCR or Northern blot (Fig. S2b-c). We cannot exclude that HPV18 circE7 is present in these cells at levels below the sensitivity of these in vitro assays.

**Figure 1.**
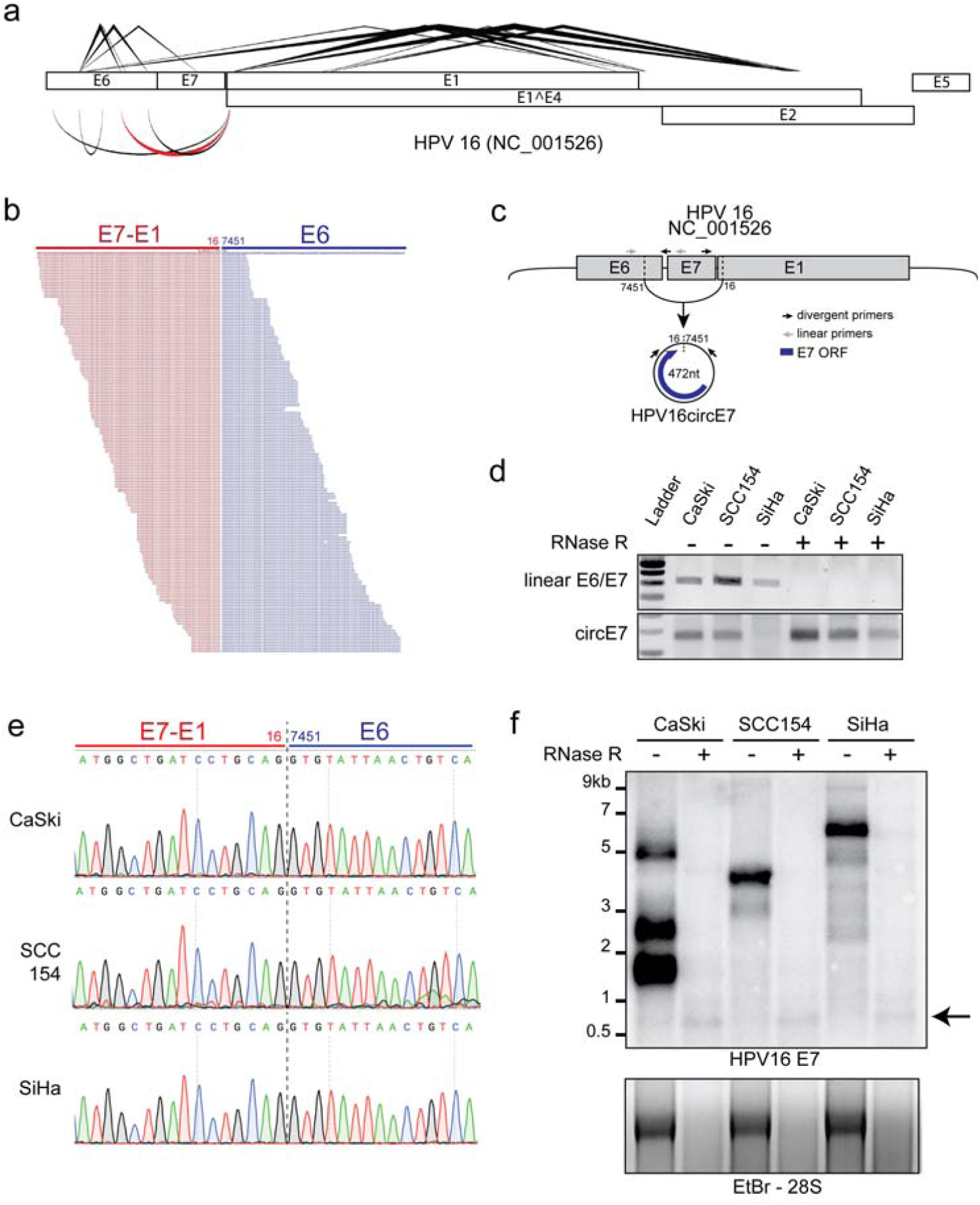
Identification of HPV circRNAs. **(a)** Diagram generated by vircircRNA summarizing the splicing events identified for HPV16 from the combined SRA datasets (Fig. S1d). Lines (top) indicate forward splicing events; arcs (bottom) indicate putative backsplicing; thickness=log_2_(read count); red highlights circE7. **(b)** Alignment of sequencing reads spanning the circE7 backsplice junction from SRS2410540. Red indicates E7-E1 sequences, and blue indicates E6 sequence. **(c)** Predicted formation and size of HPV16 circE7. Arrows indicate primers used to detect linear E6/E7 and circE7. **(d)** RT-PCR of random hexamer primed total RNA from HPV16+ cancer cell lines. 2μg of total RNA were treated with 5U of RNase R (or water for mock treatment) in the presence of RNase inhibitor for 40 min prior to RT reaction. Results are representative of 4 independent experiments. **(e)** Sanger sequencing of PCR products from (d) confirmed the presence of the expected circE7 backsplice junction without the insertion of additional nucleotides. Sequencing traces were identical for 3 independent reactions from each cell line. **(f)** Northern blot of total RNA after mock (8μg) or with RNase R treatment (20μg) from the indicated HPV16+ cell line probed with HPV16 E7. Arrow indicates RNase resistant band with E7 sequence. Ethidium Bromide staining (bottom), RNase R treatment control. Results representative of 5 independent Northerns.

CircRNAs have been reported to function by sponging miRNAs^15, 16, 17^. To determine whether circE7 might have a role as a miRNA sponge, we determined whether any miRNA binding sites existed on the transcripts. While both HPV16 HPV35 circE7 were predicted to have miRNA binding sites (Fig. S3a-b), none of the predicted miRNA binding sites were conserved between the HPV16 and HPV35 circE7 species^18^. Since circRNAs have the potential to encode peptides, we next tested whether circE7 might be translated. To facilitate the detection of circE7 translation, we generated minigene expression vectors encompassing the entire ∼1kb backspliced region of the HPV16 genome (Fig. 2a). All constructs were flanked by quaking (QKI) protein binding sites, which facilitate the circularization of RNAs^19^. Some of the constructs contained mutations of the potential start codons (no ATG) and/or a C-terminal 3xFLAG epitope tag to facilitate the detection of E7. Human embryonic kidney cells (HEK293T) were transfected with the minigenes and assayed for circRNA formation by RT-PCR using divergent primers. We detected RNase R-resistant circRNAs of the expected size from both WT and epitope tagged circE7 (Fig. 2b). When HEK293T cells were transfected with the circE7 minigenes, we were able to detect E7 protein using both HPV16 E7-specific and Flag antibodies, but not when the E7 start codons were mutated (Fig. 2c-d). To confirm that circular, rather than linear E7 RNAs, were responsible for E7 translation, we designed small interfering RNAs (siRNAs) to target sequences specific to the backsplice junction, the linear mRNA, or a region shared by both linear and circular species (Fig. 2a). RT-PCR confirmed that siRNAs against the circE7 backsplice preferentially knocked down the circular transcript; those against linear E6/E7 preferentially depleted the linear transcript; and those targeting E7 knocked down both RNAs (Fig. S3c). Notably, knockdown of circE7 or shared regions of the E7 ORF inhibited the expression of E7 protein. In contrast, siRNAs targeting the linear RNA did not strongly decrease E7 expression by Western blot (Fig. 2c). CircRNAs are predicted to be translated through a cap-independent mechanism, which can be upregulated by cell stressors, including heat shock^20, 21^. 293T cell transfected with wild-type or Flag-tagged circE7 constructs increased E7 translation, >four-fold and >two-fold, respectively, in response to a 42°C heat shock (Fig. 2d, S3d). In contrast, a linear control RNA (Flag-GFP) showed a >two-fold decrease in expression after heat shock (Fig. 2d, S3d).

**Figure 2.**
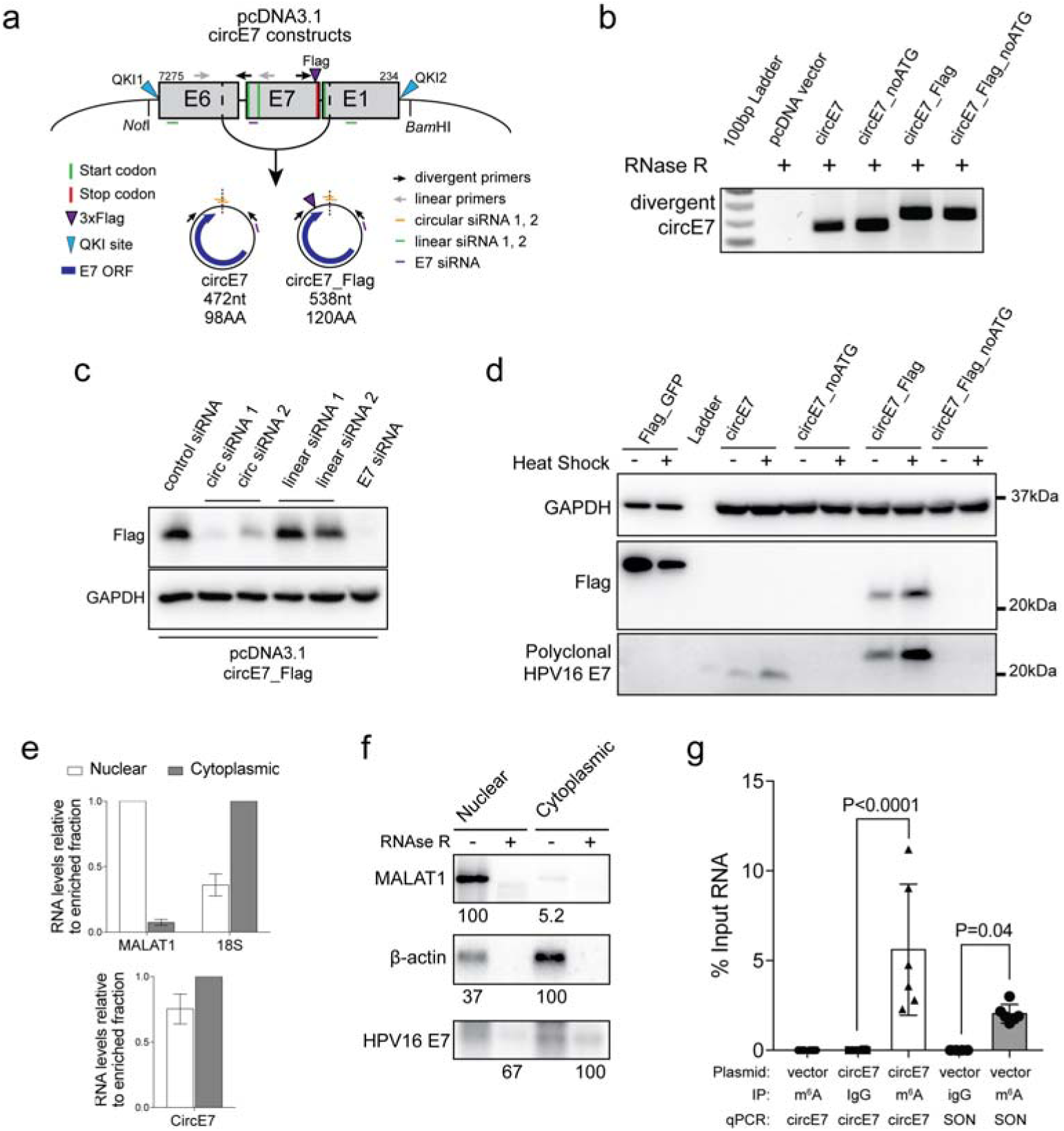
Properties of circE7. **(a)** Diagram of constructs generated to express circE7 in vitro. Map indicates location of start codons (mutated in circE7_noATG), 3xFlag (circE7_Flag), and siRNA used in subsequent experiments. **(b)** RT-PCR confirms the formation of RNase R resistant circE7 RNAs of from transfected 293T cells. Results representative of 3 independent experiments. **(c)** Western blot for Flag from 293T cells co-transfected with circE7_Flag and the indicated siRNA. GAPDH, loading control. Results representative of 4 independent transfections. **(d)** Western blots for Flag and HPV16 E7 from 293T cells transfected with the indicated circE7_Flag construct (4 μg) or a linear Flag-GFP control vector (0.4 μg). Indicated transfections were subjected to a heat shock (2 hr at 42°C, 2hr recovery). “noATG” constructs have E7 start codons mutated. Eight-fold less lysate was loaded for control Flag-GFP transfected cells. GAPDH, loading control. Results representative of 5 independent experiments. **(e)** CircE7 transfected cells were fractionated and nuclear and cytoplasmic fractions analyzed by qRT-PCR. MALAT1 and 18S (top), fractionation controls. Values normalized to the enriched fraction. Results are representative of 3 independent fractionations. **(f)** Northern blot of total RNA (4 μg) with mock or with RNase R treatment from fractions of 293T cells confirms that circE7 is enriched in the cytoplasm and is RNase R resistant. MALAT1 and β-actin, fractionation controls. Numbers (bottom) indicate quantitation of band density normalized to the enriched fraction. Results are representative of 3 independent blots. **(g)** RT-PCR of RNA IP (m^6^A or IgG control) after transfection with the indicated plasmid (24hr) (n=6 biological replicates from 3 transfections). SON, m^6^A RNA IP control. Data are shown as mean±s.d. *P* values (indicated above relevant comparisons) were calculated with one-way analysis of variance (ANOVA) with Holm-Sidak tests (g).

To determine the subcellular localization of circE7, cells were fractionated and nuclear and cytoplasmic fractions confirmed with MALAT1 and 18S or β-actin, respectively. Consistent with the behavior of other translated circRNAs^20, 22, 23, 24^, RT-PCR confirmed that the majority of circE7 (∼60%) localized to the cytoplasm (Fig. 2e). Northern blots confirmed the cytoplasmic enrichment of circE7 (Fig. 2f). Cap-independent translation of circRNAs has been reported to require N^6^-methyladenosine (m^6^A) modifications in the UTR^20^. Because HPV16 circE7 possessed multiple potential m^6^A consensus sites (DRACH) (Fig. S3d), we performed m^6^A RNA immunoprecipitation (IP) experiments. Antibodies against m^6^A, but not an IgG control, pulled down circE7 even more efficiently than SON, a control mRNA containing multiple m^6^A sites and previously confirmed to be methylated^20^. We constructed a circE7 mutant in which potential m^6^A motifs in the UTR were mutated (circE7_noDRACH) (Fig. S3e). Unexpectedly, this construct dramatically decreased the abundance of circE7, but not its linear E6/E7 counterpart (Fig. S3f). Mutation of the potential m^6^A motifs also strongly inhibited E7 expression, further confirming the critical role for the circular, rather than the linear, RNA in E7 translation (Fig. S3g). Thus, circE7 is m^6^A-modified, enriched in the cytoplasm, and capable of generating the E7 oncoprotein in a heat shock regulated manner.

The functions of most circRNA remain ambiguous. In particular, the possible functions of virally-encoded circRNAs and those purported to encode for proteins remain particularly poorly characterized. To determine the biological functions of circE7, we specifically depleted circE7 in CaSki cells using two doxycycline-inducible short hairpin RNAs targeting the circE7 backsplice junction (circE7 sh1/2). After lentiviral transduction of the circE7 shRNA-expressing plasmid, we confirmed the specificity of the circE7 shRNA by qRT-PCR. After doxycycline induction, both circE7 shRNA resulted in a significant reduction of the circE7 as assessed both by RT-PCR and Northern blot (Fig. 3a-b). Importantly, we did not note a significant reduction of the linear E6/E7 sequences or the previously described E6*I transcript (Fig. S4a-b). Unexpectedly, both qRT-PCR and Northern blot suggested that knockdown of circE7 actually caused an increase in linear HPV16 E6/E7 transcripts (Fig. S4a-b). Next, we tested whether loss of circE7 would impact levels of E7 protein in CaSki cells. Induction of circE7 shRNA 1/2 decreased levels of endogenous E7 protein by >two-fold (Fig. 3c, S4c), demonstrating that circE7 is required for optimal E7 expression in CaSki cells. Consistent with E7’s established role in transformation, depletion of circE7 resulted in decreased cell proliferation as measured by both cell number and MTT assay (Fig. 3d; S4d-e). CaSki cells expressing circE7 shRNA showed significantly decreased entry into S phase as measured by BrdU incorporation (Fig. 3e, S4f) consistent with a critical role for E7 in overriding Rb’s function in regulating cell cycle progression^25^. Induction of circE7 sh1/2 also significantly inhibited the ability of CaSki cells to form colonies in soft agar (Fig. 3f). We conclude that circE7 is essential for E7 expression and the transformed growth of CaSki cervical carcinoma cells.

**Figure 3.**
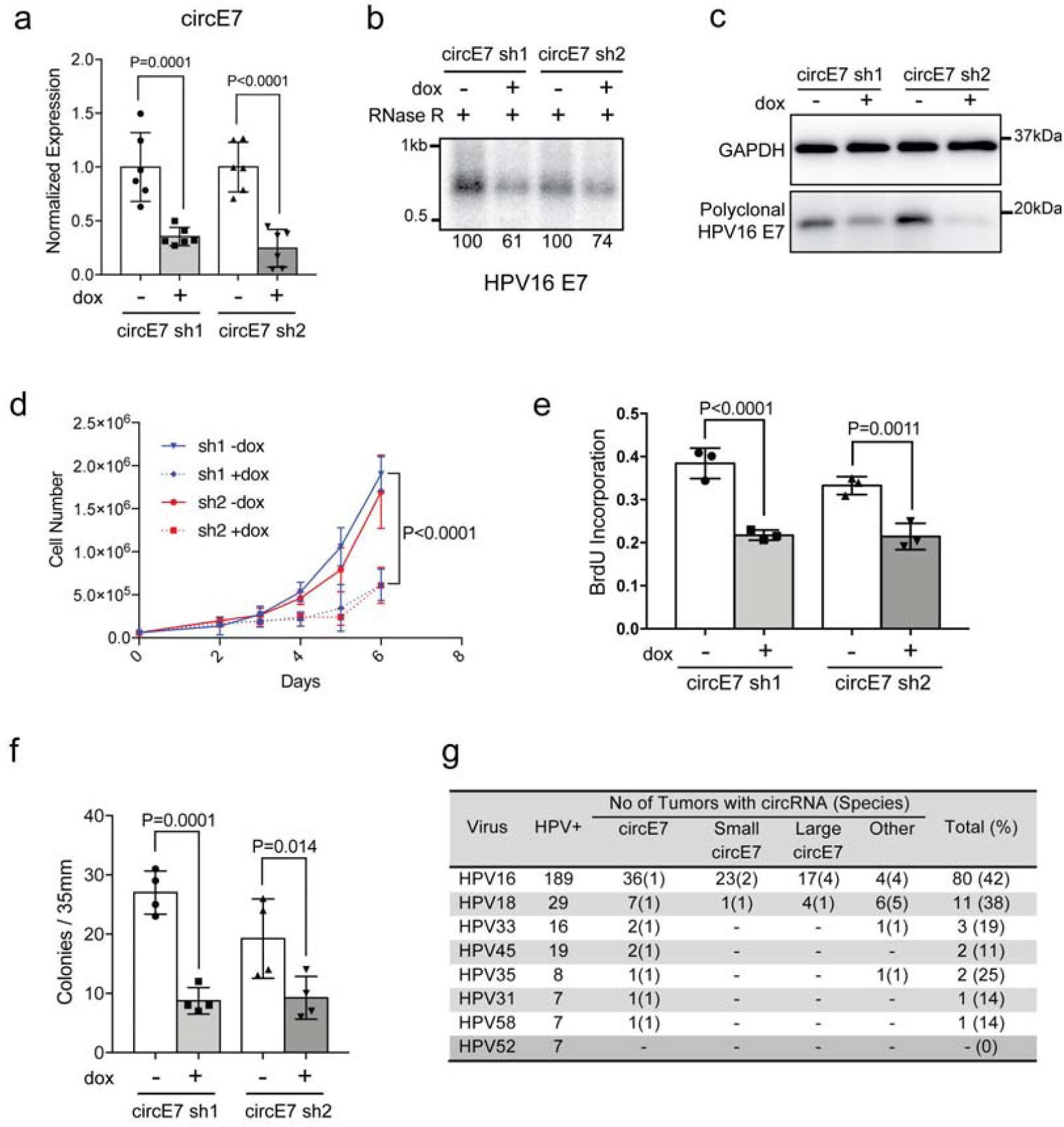
Biological function of circE7. **(a)** CaSki cells were lentivirally transduced with doxycycline (dox)-inducible hairpins specific for the circE7 backsplice junction (circE7 sh1/2). qRT-PCR of random hexamer primed total RNA revealed that circE7 sh1/2 resulted in significant decreases of circE7 levels. (n=3 independent experiments, run in duplicate) **(b)** Northern blot of RNase R treated total RNA (30 μg) from CaSki cells with or without circE7 sh1/2 induction (2 days). Numbers (bottom) indicate quantitation of band density normalized to the uninduced control. **(c)** CircE7 sh1/2 induction (3 days) resulted in the decreased expression of endogenous E7 expression. Western blots are representative of 3 independent experiments. **(d)** 6.0×10^4^ CaSki cells were seeded in triplicate in six-well plates at day 0 and absolute cell number quantitated daily after day 2. CircE7 sh1/2 induction resulted in significantly slower growth of CaSki cells after day 4. Similar results were obtained in 3 independent experiments. **(e)** CaSki cells with or without circE7 sh1/2 induction (1 day) were plated in chamber slides and labeled with BrdU (10μM for 1.5 hrs). Cells were stained with αBrdU and DAPI and scored as % of DAPI+ cells. **(f)** 1.0×10^4^ CaSki circE7 sh1/2 cells with or without induction (1 day) were seeded in triplicate in soft agar with or without dox (14 days). Average colonies per 35mm. n=4 independent transfections. **(g)** TCGA RNA-Seq data (CESC, HNSC) was analyzed with vircircRNA and backsplices with ≥2 reads were tabulated. Data are shown as mean±s.d. *P* values (indicated above relevant comparisons) were calculated with two-tailed *t* test (d) and one-way analysis of variance (ANOVA) with Holm-Sidak tests (a,e,f).

Given the critical role of circE7 in maintaining transformation in CaSki cells, we tested whether circE7 might be more broadly relevant in human cancers. We analyzed TCGA RNA-seq data from cervical squamous cell carcinoma and endocervical adenocarcinoma (CESC) and head and neck squamous cell carcinoma (HNSC) using the vircircRNA pipeline and identified more than 100 patient tumors with at least two reads identifying the same HPV backsplice junction. To ensure the specificity of the identified HPV reads, a control analysis of kidney renal clear cell carcinoma (KIRC), which has been reported to be free of HPV transcripts^26^, yielded no HPV sequence reads or backsplice junctions using identical search parameters. Backsplice junctions were detected from multiple high-risk HPV, with HPV16 being the most abundant species (Fig. 3g). Consistent with our preliminary pipeline analysis, multiple species of circE7 were the most abundant type of backsplice identified in all high-risk HPV species (>95% of total species). In contrast to our in vitro analyses, HPV18+ patient tumors also appeared to possess frequent backsplices consistent with HPV18 circE7.

## Discussion

We describe the functional activity of a viral, protein-coding circRNA. Despite its low abundance relative to linear HPV transcripts, circE7 appears to play a critical role in HPV16’s ability to transform CaSki cells. Post-transcriptional modifications (e.g. m^6^A) and the cytoplasmic localization of circE7 may explain, in part, the striking contribution of circE7 to E7 oncoprotein expression. While HPV16 circE7 could readily be detected through both in vitro assays from cancer cell lines and in TCGA RNA-Seq data, HPV18 circE7 could only be identified in TCGA RNA-seq datasets. While the lower sensitivity of in vitro assays could explain this discrepancy, we speculate that the absence of circE7 in HPV18+ cancer cells actually highlights differences in the regulation of circRNA formation. Previous studies have suggested that both cis (locus specific) and trans (host cell specific) factors determine the rate of backsplicing events^19^. For example, transcripts with a strong tendency to circularize would be abundant regardless of the host tissue type or differentiation state, whereas more weakly circularizing transcripts would only circularize with the tissue specific expression of RNA binding proteins. Similarly, HPV16 circE7 may have cis properties that promote constitutive circularization while HPV18 might only promote circularization under specific growth or differentiation conditions.

HPVs modulate their transcription, polyadenylation, and linear splicing to respond to the differentiation state of the epithelial cells in which they reside. The formation of circRNAs adds another layer of complexity to how HPVs could regulate infection and immune evasion. In the absence of E6, the expression of the E7 oncoprotein upregulates p53 activity and promotes apoptosis^25, 27^. Thus, the low translational activity of circE7, in addition to the established stability of circRNAs relative to linear mRNAs, might make them particularly well-suited to promote the fitness of infected cells during latency. Our studies also implicate m^6^A in the regulation of circE7. While a previous study implicated RNA methylation specifically in circRNA translation^20^, the mutation of the abundant m^6^A-consensus motifs in the UTR of circE7 dramatically decreased the efficiency of backsplicing in our assays. Although m^6^A-motifs do not appear to be essential for splicing^28^, we speculate that m^6^A deposition on the nascent circE7 RNA may somehow coordinately regulate backsplicing and translation. While the m^6^A motif sites do not correspond to the binding sites for factors known to regulate HPV splicing^29^, we have not excluded the possibility that the mutations might also directly impact circE7 backsplicing by altering the binding of canonical splicing factors.

While our studies focused on circE7, we speculate that other viral and protein-coding circRNAs will also ultimately be shown to have biologically relevant properties. The functional activity of circE7 also suggests important avenues for additional investigation. The compact size of circE7 may be useful as a template for understanding how backsplicing is regulated. Such studies may also yield even novel insights on how HPVs regulate infection, latency, and tumorigenesis. The detection of circE7 may also have clinical implications. Given the stability of circRNA and the importance of the E7 oncoprotein in tumorgenesis, it will be worthwhile determining whether circE7 would be useful as a diagnostic test. While the utility of high-risk HPV testing for cancer screening is well established, it will be interesting to determine whether the specific presence of the circE7 backsplice junction has any prognostic significance. As technologies to deliver RNAs for therapeutic benefit mature, protein expressing circRNAs, like circE7, may serve as a backbone for the design and optimization of protein-expressing circRNAs.

## Acknowledgements

We thank K. Ariizumi, J. Chung, J. Min, and J. Shay for use of hybridization equipment; A. Brown and N. Conrad for use of the Typhoon PhosphorImager and assistance with cell fractionation; and D. Castrillon for cervical carcinoma cells. We thank N. Conrad, T. de Lange, J. Shay, P. Tsai, and K. Yancey for critically reviewing the manuscript. This work was supported by grants from the UT Southwestern Cary Council, Burroughs Wellcome Fund (1010978), ACS (RSG-18-058-01), and NIAMS (R01AR072655) to R.W.

## Author contributions

E.L., J.K., and R.W. performed experiments related to Fig. 1. J.Z., E.L., and R.W. performed experiments related to Fig. 2 with assistance from R.Y. J.Z., J.K., and R.W. peformed experiments related to Fig. 3 with assistance from B.H. and E.L. J.K. performed all bioinformatic analyses with input from M.K. and X.Z. R.W. conceived the study, designed the experiments, and wrote the paper with input from all co-authors. UT Southwestern has filed a patent on this technology with E.E.L., J.Z., and R.C.W. named as inventors.

---

**Figure S1.**
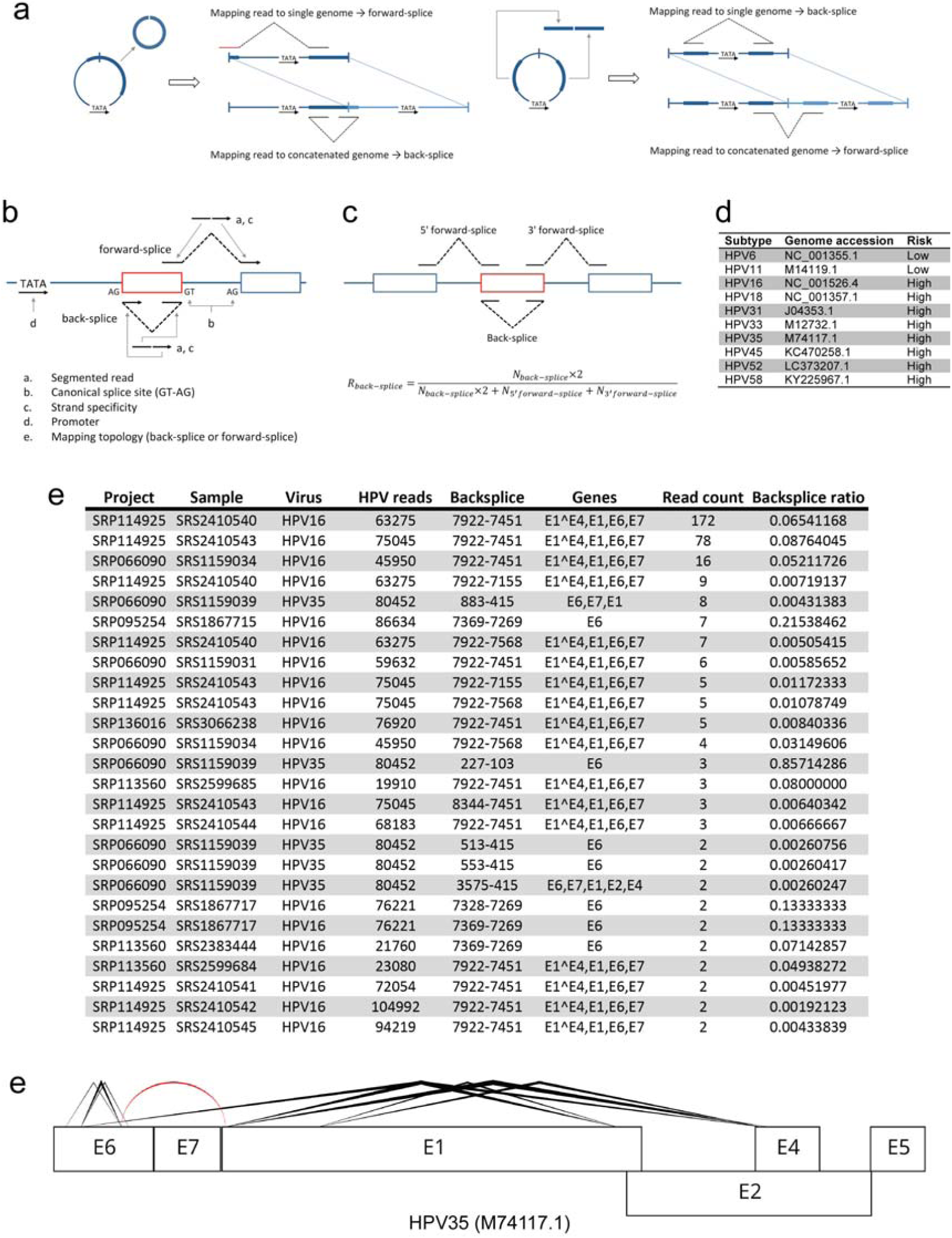
Identification of HPV circRNA. **(a)** Schematic demonstrating the rationale for concatenation of linearized viral genomes in the vircircRNA pipeline. This strategy allows for the accurate identification of both back and forward splices. **(b)** Criteria employed by vircircRNA to identify splice sites. **(c)** Schematic and formula used to calculate backsplice ratios. **(d)** Type, genome, and risk category of HPV genomes used in vircircRNA analysis. **(e)** Summary of HPV circRNAs identified in SRA datasets including location, genes contained in putative circRNA, read count, and backsplice ratio. **(e)** Diagram generated by vircircRNA summarizing splicing events identified for HPV35. Lines indicate linear splicing; arcs indicate circular splicing; thickness=log_2_(read count); red highlights circE7.

**Figure S2.**
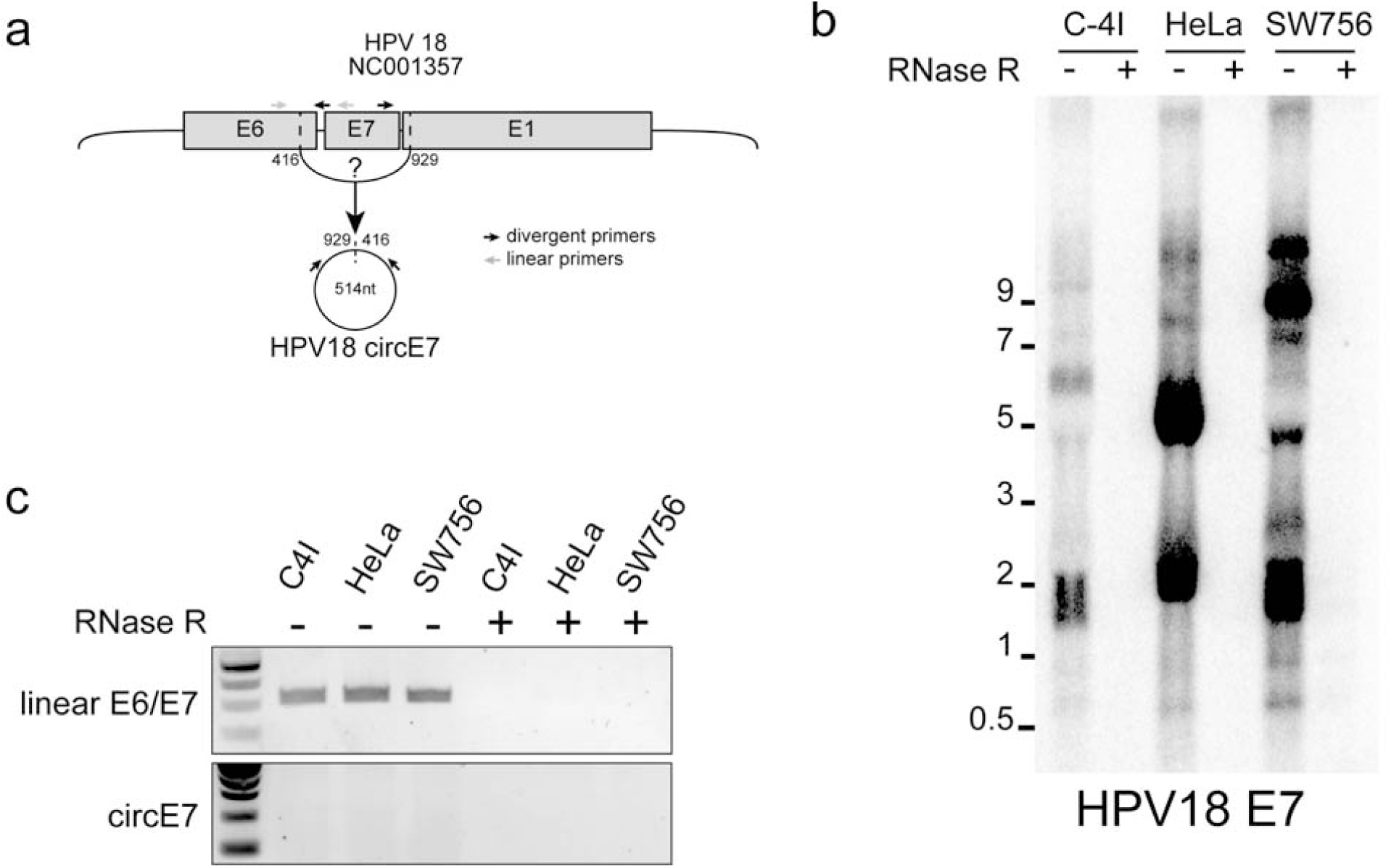
Absence of HPV18 circE7 from HPV18+ cell lines. **(a)** Predicted size and formation of putative HPV18 circE7. Splice sites were derived from the analogous splice sites in HPV16. Arrows indicate primers used to detect linear E6/E7 and circE7. **(b)** RT-PCR of random hexamer primed total RNA with or without RNase R treatment reveals loss of linear mRNA. A product consistent with HPV18 circE7 was not detected. **(c)** Northern blot using HPV18 E7 as a probe identifies did not identify RNase R resistant bands in HPV18+ cell lines. Total RNA after mock treatment (8μg) or after RNase R treatment (20μg) from the indicated HPV18+ cell line.

**Figure S3.**
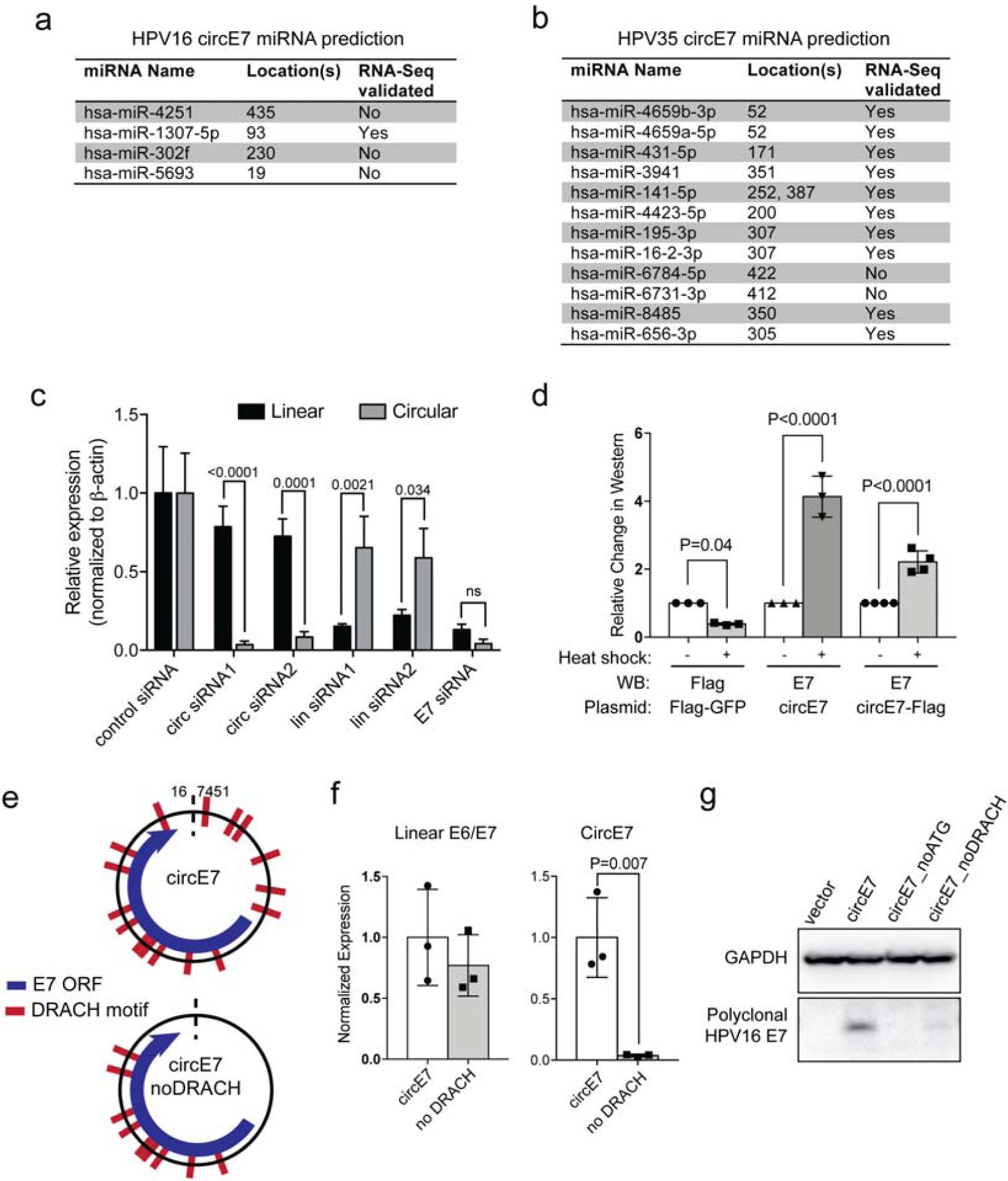
Properties of circE7. **(a-b)** Predicted miRNA binding sites in HPV16 circE7 (a) and HPV35 circE7 (b) (mirDB). No miRNAs are predicted to be conserved between the circE7. **(c)** Quantitative RT-PCR from pcDNA3.1-circE7_Flag transfected 293T cells confirms that circ siRNAs significantly deplete circE7 compared to the linear E6/E7; lin siRNAs significantly deplete the linear mRNA; and E7 siRNA depletes both isoforms. **(d)** Quantitation of band density of Western blots for Flag and HPV16 E7 from 293T cells transfected with the indicated plasmid (Fig. 2D). Values normalized to the control (no heat shock) cells which were not subjected to heat shock (2 hr at 42°C, 2hr recovery). **(e)** Schematic of the DRACH consensus motifs for METTL3/14 and the sites mutated in the circE7_noDRACH construct. **(f)** qRT-PCR of circE7 or circE7_noDRACH transfected cells. Loss of UTR DRACH motifs in circE7 results in a significant decrease in the abundance of circE7, but not linear E6/E7. n=3 independent experiments, run in duplicate. **(g)** Mutation of DRACH motifs and loss of circE7 markedly decreases expression of the E7 oncoprotein in transfected 293T cells. Data are shown as mean±s.d. *P* values (indicated above relevant comparisons) were calculated with two-tailed *t* test (f) and one-way analysis of variance (ANOVA) with Holm-Sidak tests (c,d).

**Figure S4.**
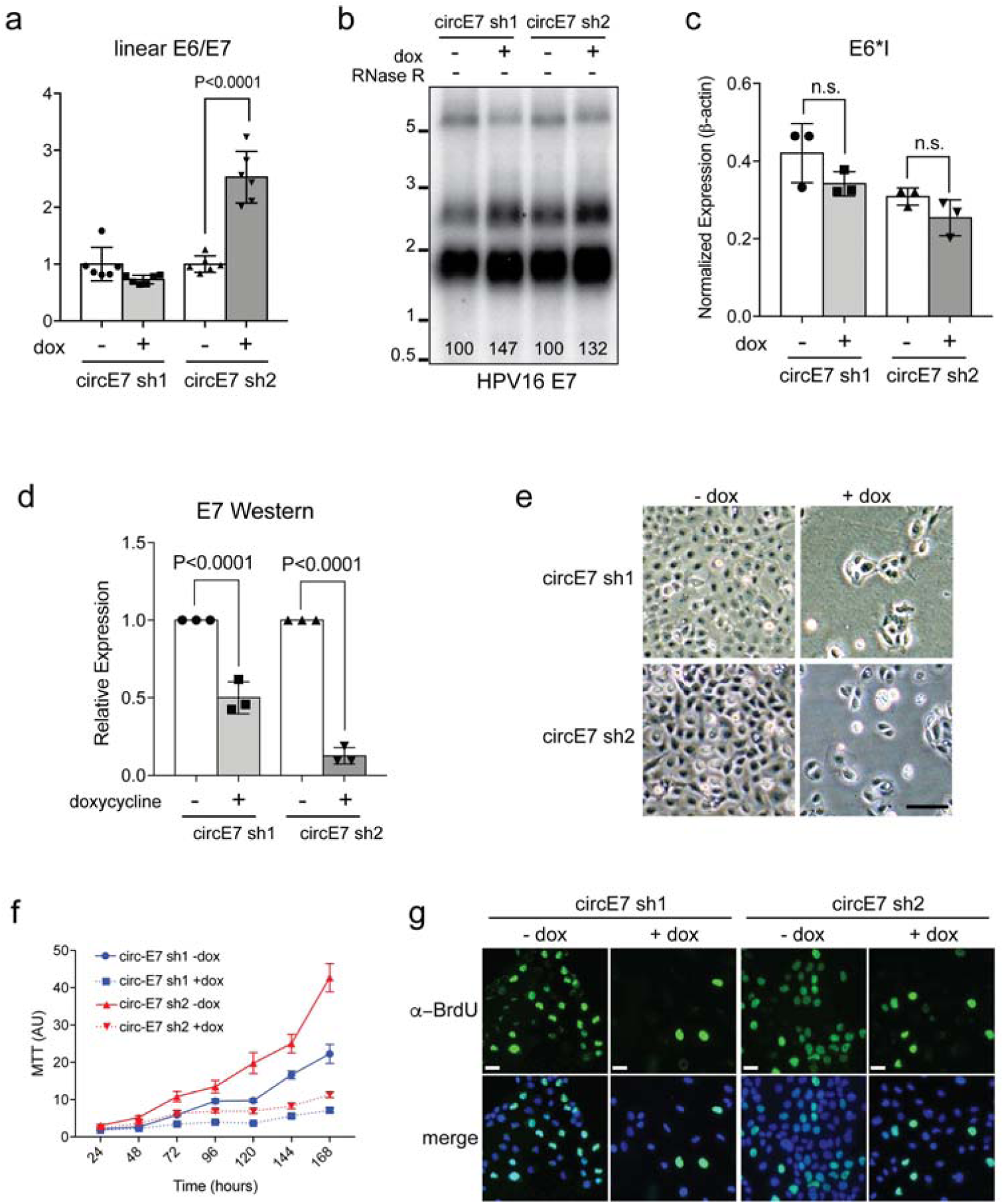
Biological functions of circE7. **(a)** RT-PCR for linear E6/E7 in CaSki with or with induction of circE7 sh1/2. Induction of circE7 sh2 results in a significant *increase* of linear E6/E7 transcripts. (n= 3 independent experiments). **(b)** Northern blot of total RNA (4 μg) from CaSki cells with or without circE7 sh1/2 induction (2 days). Numbers (bottom) indicate quantitation of band density normalized to the uninduced control. **(c)** qRT-PCR for E6*I transcript in CaSki with or with induction of circE7 sh1/2. Induction of circE7 sh1/2 does not significantly change E6*I transcript abundance. **(d)** Quantitation of band density of Western blots for HPV16 E7 from CaSki cells with or without circE7 sh1/2 induction. Values normalized to the uninduced condition. **(e)** Differential interference contrast (DIC) images of CaSki with and without dox induction of circE7 sh1/2 (5 days). Bar, 50μm. **(f)** MTT assay of CaSki circE7 sh1/2 cells with and without doxy induction. **(g)** Representative images of BrdU and DAPI staining from CaSki with and without dox induction circE7 sh1/2. Bar, 10μm.

## Materials and Methods

### De novo circular RNA detection from circular viral genomes

To detect and display circular RNA from RNA sequencing (RNA-seq) data of viruses with circular genomes, a custom pipeline named vircircRNA was developed (https://github.com/jiwoongbio/vircircRNA). Because the arbitrary linearization of circular genomes causes confusion in distinguishing between back and forward splices, two genomes were concatenated and used as the reference sequence for read mapping. RNA-seq reads were aligned using Burrows-Wheeler Aligner (BWA, v0.7.15)^1^ with specific options, “-T 19” to reduce minimum score to output and “-Y” to use soft clipping for supplementary alignments. Reads mapped to different positions with soft clipping were extracted as candidate segmented reads by splicing. Canonical GT-AG splice donor-acceptor motifs were used to exclude non-splice reads and define splice breakpoints. Information on strand specificity of sequencing and annotation of genes and promoters were included if available. Splice reads mapped in chiastic order were defined as back-splice reads from circular RNAs. The back-splice junction ratio was calculated by employing the equation:

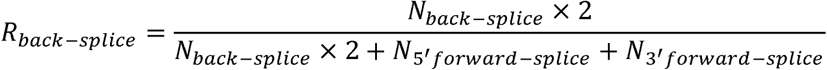

where 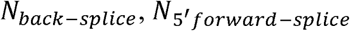 and 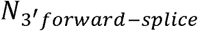 are the numbers of back-splice, 5′ and 3′ normal-splice junction reads.

### Circular RNA detection from public and TCGA RNA-seq data

For selected HPV, the genome sequences and the annotations were downloaded from National Center for Biotechnology Information (NCBI) nucleotide database. We used the keywords of “HPV” and “human papillomavirus” to search public sequencing data of HPV from NCBI Sequence Read Archive (SRA) and found 154 RNA-seq data accessions of cDNA library from 12 projects. For the TCGA data, we downloaded raw sequencing data from cervical squamous cell carcinoma and endocervical adenocarcinoma (CESC, 309 samples), head and neck squamous cell carcinoma (HNSC, 566 samples), and kidney renal clear cell carcinoma (KIRC, 618 samples) from The Cancer Genome Atlas (TCGA). To avoid false mapping, reads (50-51bp) were mapped onto both HPV genomes and human transcript sequences downloaded from Ensembl (release 92, GRCh38) (https://www.ensembl.org/). Only reads with alignment scores on HPV genomes greater than those on human transcripts were kept for further analysis. HPV-positive samples were defined by ≥10,000 reads mapped on a HPV genome and a mapping rate on the genome ≥0.00001. The survival analysis was performed using R packages, survival and survminer.

### Cell Culture

C-4I cells were cultured in Waymouth’s and 10% FBS. CaSki (ATCC) cells were cultured in RPMI and 10% FBS. HeLa (ATCC), SW756, and SiHa cells were cultured in DMEM and 10% FBS. UPCI-SCC154 (ATCC) cells were cultured on Collagen I coated plates in DMEM/F-12 (Gibco, 11320082) supplemented with GlutaMAX (Gibco, 35050061), 10% FBS, 0.1 μg/ml hydrocortisone (Sigma, H0888), and 10 ng/ml EGF (Invitrogen, 10450-013).

### Northern blots

Total RNA from indicated cells was extracted by Qiagen RNeasy. Similar results were obtained using TRIzol (Invitrogen). For cancer cell lines, 20-30µg of total RNA was treated with RNase R or 8µg of total RNA used for mock treatment. Similar results were obtained with using rRNA depleted samples (NEBNext) with 4ug of depleted RNA for each sample. For fractionated RNA sample, 4µg of total RNA per fraction was used. RNA samples were heated in 2x formamide sample buffer and were electrophoresed in 1.5-2.0% formaldehyde-agarose gels in MOPS buffer and transferred onto Hybond N+ membranes (Amersham) with 10xSSC. Probe fragments were generated by PCR using the primers indicated in Table S2. PCR products were gel purified (Machery Nagel) and used to generate α-^32^P dCTP-labeled probes using the Random Prime Labeling Kit (Takara/Clontech). Probes were purified over nucleotide purification columns (Zymo Research) prior to hybridization. Hybridization was performed in PerfectHyb buffer (Sigma) at 65°C overnight and washed according to Hybond’s instructions. Blots were developed on PhosphorImager screens and developed on a Typhoon Imager. Blots were quantited with ImageQuant.

### Circular RNA Transient Expression Constructs and Transient Expression

Circular E7 RNA cDNA were synthesized as gBLOCK fragments (IDT), PCR amplified, and cloned into pCDNA3.1- vector by *Not*I/*Bam*HI restriction sites. Positive clones were verified by Sanger Sequencing (Genewiz). Constructs were transiently transfected into LentiX-293T (Clontech) cells by Lipofectamine 3000 including the P3000 reagent (ThermoFisher). At 48-72hr post-transfection, cells were harvested for fractionation, end-point PCR, RT-PCR, or Western blot.

### Western blotting

Whole-cell protein extracts were separated on SDS-PAGE gels, transferred to PVDF membranes and probed with the following primary antibodies: anti-FLAG HRP (1:300, Sigma, A8592), anti-GAPDH (1:1200, Santa-Cruz, sc-32233), and anti-HPV16 E7 (1:1000, GeneTex, GTX133411). Membranes were then incubated with the appropriate HRP-conjugated secondary antibody (1:000, anti-mouse IgG, Cell Signaling Technologies, 7076P2; 1:1000, anti-rabbit IgG, ThermoFisher, G21234) and developed with an ECL system (Perkin Elmer, NEL104001). CaSki cells were harvested and lysed in RIPA buffer (50mM Tris pH 8.0, 150 mM NaCl, 1% NP-40, 0.1% SDS, 0.5% Sodium Deoxycholate) in presence of 1x protease/phosphatase inhibitor. Cells were incubated on ice with RIPA buffer for 10-15min. Whole cell lysate was then cleared by centrifugation at 12,000Xg for 15 min. The cleared supernatant was transferred to new tubes, before added Laemmli sampling buffer and boiled for at 95°C for 10min.

### RNA isolation, RNase R treatment, cDNA synthesis

Total RNA was extracted from cells using the RNeasy Mini Kit (Qiagen, 74104) and RNase R digestion was performed as previously described^2^. Similar results were obtained with TRIzol reagent (Sigma, T9424). 2µg of total RNA was incubated with 5U RNase R (Lucigen, RNR07250), 10U murine ribonuclease inhibitor (New England Biolabs, M0314S), 0.5U DNase (Qiagen, 79254), and 1X RNase R buffer for 40 min at 37°C and then placed on ice. Water was substituted for RNase R in mock reactions. 1µl of 1mM EDTA and 1µl random hexamer (100µM) were added to the unpurified RNase R digested products and denatured at 65°C for 5min. Samples were then placed on ice and reverse transcribed using the Superscript IV RT system (ThermoFisher, 18091050). For standard cDNA synthesis (no RNase R), 1µg of total RNA was then reverse transcribed using Bio-Rad iScript cDNA synthesis kit according to manufacturer’s instruction.

### End-point PCR and qRT-PCR

End-point PCR was performed with SapphireAmp (Takara, RR350B) by denaturation at 95°C for 5min, followed by 23 cycles at 95°C 1min, 62°C 30sec, 72°C 1min, and a final extension step at 72°C for 7min for linear products. Cycling conditions for circRNA: were as follows: 95°C 5min, followed by 40 cycles of 95°C 1min, 62°C 1min, 72°C 2min, and a final elongation step at 72°C for 10min. Cycling conditions for linear mRNA: 95°C 5min, followed by 23 cycles of 62°C 30sec, 72°C 1min, 72°C 7min, 23 cycles. For endpoint PCR, cDNA templates were diluted 1:4 in water for linear PCR reactions. For qRT-PCR, cDNA products were diluted 1:20, 2µL of diluted DNA sample was used as template for real time PCR analysis with PowerUp SYBR Green (Applied Biosystems, A25779). Data were normalized by β-actin as a loading control and then presented as relative normalized expression level.

### Nuclear and cytoplasmic fractionation

Cell fractionation was performed as previously described with minor modifications3. Cells grown in 35mm dishes were trypsinized and centrifuged at 500g for 3min at 4°C. Pelleted cells were washed once with ice-cold PBS, centrifuged again, and resuspended in 250µl of ice-cold Buffer I [0.5% Triton X-100, 0.32M sucrose, 3mM CaCl_2_, 2mM MgCl_2_, 0.1mM EDTA, 10mM Tris (pH 8.0), 50mM NaCl, 1mM DTT, 0.04U/µl RNase inhibitor (ThermoFisher, 18091050)]. After a 15 min incubation on ice, cells were centrifuged at 500g for 5min at 4°C. The supernatant was collected for the cytoplasmic fraction and the pellet was resuspended in 250µl of Buffer I for the nuclear fraction. RNA was extracted from the cytoplasmic and nuclear fractions using the RNeasy Mini Kit (Qiagen, 74104) and reverse transcribed with Superscript IV RT system (ThermoFisher, 18091050) using random hexamers.

### m^6^A RNA Immunoprecipitation Assay

pCDNA3.1-empty vector or circE7 expression constructs were transiently transfected into LentiX-293T cells in triplicates. Total RNA was harvested 48-72hr post-transfection. 5µg total RNA was immunoprecipitated (IP) by 1µg m6A polyclonal antibody (Synaptic Systems#202003). Immunoprecipitated RNA was then purified over QIAgen RNeasy columns. Purified immunoprecipitated RNA, along with 10% input RNA were then reverse transcribed by Bio-Rad iScript DNA synthesis kit and analyzed by real time PCR. Percentages of m6A modified RNA for both circE7 were calculated based on the input reading.

### Doxycycline inducible shRNA Construct and Lentiviral packaging

Short hairpin RNA (shRNA) sequences specifically targeting circE7 back-splice junction were designed from Invitrogen BLOCK-ItTM RNAi Designer (https://rnaidesigner.thermofisher.com/rnaiexpress/). Forward and reverse oligos were annealed in NEB buffer 2, and then phosphorylated and ligated into EZ-Tet-pLKO-Blasticidin vector^4^ (Addgene #85973) using *Nhe*I/*Eco*RI. To package lentiviruses, LentiX-293T cells were plated at ∼70%-80% confluence 12-16hrs before transfection. Transfer plasmid, pMD2.G and psPAX2 were then co-transfected into LentiX-293T cells by Lipofectamine 3000 at molar ratio of 1:1:1. Viruses were harvested at 48hr and 72hr post-transfection. For lentiviral transduction, CaSki cells were plated into 6 well plate at 50% confluence without any antibiotics. Viral supernatant was then added to the cell in presence of polybrene at final concentration of 8µg/mL. 48hrs after transduction, the cells were then exposed to blasticidin selection at final concentration of 10µg/mL for the first day and then increased to 15µg/mL on the second day of selection. Cells were selected for ≥7days before the antibiotic concentration was decreased to 5µg/mL for maintenance of stable cell lines. For shRNA expression induction, doxycycline was added to culture medium at final concentration of 1µg/mL for at least three consecutive days and cells were then subjected to analyses.

### Cell proliferation and MTT cell growth assay

For the cell proliferation assay, 60,000 CaSki cells were plated on Day 0 in quadruplicate RPMI with 10%FBS (doxycycline free) with or without 1µg/mL doxycycline. Cells were counted daily after Day 2. For MTT based cell growth rate analysis, 1000 CaSki cells stably transduced with two shRNA constructs were plated into each well of 96 well plates in triplicates for at least 8 days without doxycycline. The first time point was done 24hrs after plating, serving as a reference point. After 24 hrs, doxycycline was added to the rest of plates. The MTT assay was performed daily for 7 days. All the readings from the same well were normalized by the reference point. MTT (Invitrogen#M6494) reactions were performed as recommended by manufacturer.

### BrdU Incorporation Assay

CaSki cells stably transduced with circE7 shRNA1/2 were induced by 1µg/mL doxycycline for at least three days. Control cells (no dox) and induced cells were then plated into Nunc 4-well chamber slides (ThermoFisher 154453) in triplicate. Cells were labeled with 10µM BrdU for 1.5hrs. Cells were then fixed by 4% paraformaldehyde at RT for 10mins, followed by 0.1% Triton X-100 permeabilization and 1.5N HCl DNA hydrolysis. Cells were then probed by O/N incubation with BrdU antibody (1:400, BD BioScience) and stained with Alexa 488 conjugated secondary antibody (ThermoFisher). Samples were then subjected to DAPI nuclear counterstain (Vector Labs). BrdU positive cells were then quantified and data were presented as percentage BrdU positive cells.

### Soft Agar Colony Formation Assay

CaSki cells stably transduced with the two doxycycline inducible shRNA constructs were induced by 1µg/mL doxycycline for 3 days. 0.5% agar, DMEM, 10%FBS was plated as base layer in 6-well plate (35mm). 10,000 cells of each group were plated in 0.3% agar-DMEM, 10%FBS on top of the base layer in triplicates. Cells were fed with 500µL RPMI with 10%FBS medium twice a week. Doxycycline (1µg/mL) was included in all media for induced cells. Cells were then allowed to grow for 14 days before quantification. Colonies larger than 100μm were scored.

